# Fungal endophytes of cactus (*Stenocereus* spp.) as a potential alternative to alleviate drought stress in juveniles of *Theobroma cacao* L. ICS95

**DOI:** 10.1101/2025.09.16.676698

**Authors:** Karen Sofía Trujillo-Ortigoza, Angelis Marbello-Santrich, Fermín Rada, Marcela Guevara-Suarez, Silvia Restrepo

## Abstract

*Theobroma cacao*, one of Colombia’s most economically and socially significant crops, faces productivity challenges due to drought. This stress can reduce growth, leaf area, and stomatal conductance (Ks), and generate reactive oxygen species (ROS). Therefore, exploring solutions to enhance drought tolerance is crucial. This study aimed to evaluate the use of fungal root endophytes from *Stenocereus* spp. to induce drought tolerance in *T. cacao* genotype ICS95. *In vitro* drought tolerance screening identified five fungal isolates that exhibited the highest biomass production and less than 20% biomass loss under drought compared to non-drought conditions. The soil of juvenile *T. cacao* plants was inoculated with these isolates, and physiological and morphological parameters were assessed, including leaf water potential (Ψ_L_), stomatal conductance (Ks), proline content, and growth. The results showed a significant decrease in Ψ_L_ and Ks in juveniles under drought stress, which was observed across all five fungal isolates tested. However, juveniles inoculated with *Phoma* sp. exhibited less negative Ψ_L_ and lower Ks than non-inoculated controls, suggesting that this fungus may be a potential inducer of drought tolerance in *T. cacao* ICS95. One intriguing result was that plants inoculated with this fungus accumulated less proline during the drought treatment. Under non-drought conditions, juveniles inoculated with *Acrophialophora* sp., *Ectophoma* sp., *Fusarium* sp., and *Phoma* sp. exhibited an increase in mean leaf area. These findings suggest that fungal endophytes associated with *Stenocereus* spp. could provide a potential alternative for alleviating drought stress and may also mediate growth promotion under non-drought conditions in cacao.

**Importance:** *Theobroma cacao* is among the world’s most valuable crops, yet its productivity is increasingly threatened by fluctuating rainfall and prolonged drought. Identifying sustainable strategies to mitigate these impacts is therefore critical. Xerophilic plants, such as *Stenocereus* spp., harbor diverse fungal endophytes adapted to arid environments, representing a promising source of microorganisms capable of enhancing stress tolerance in commercial crops. Our study demonstrates that cactus-derived endophytes could improve drought resilience in juvenile cacao by modulating physiological responses such as stomatal conductance and leaf water potential. Furthermore, under favorable conditions, some endophytes could promote growth by increasing leaf area compared to non-inoculated plants. These findings underscore the potential of fungal endophytes from arid ecosystems as biotechnological tools for sustainable cacao production, offering an environmentally friendly alternative to mitigate drought stress while enhancing plant performance.

## Introduction

*Theobroma cacao* L., commonly known as cacao, is a native crop of the Amazon basin. This crop plays a significant commercial role in Africa, Central America, and South America (1) (2). Cocoa beans are used to produce chocolate, pharmaceutical formulations, cosmetic products, and alcoholic beverages (1–3), making the crop a valuable commodity across various industries.

Colombia is one of the largest producers of *T. cacao,* renowned for producing fine, aromatic, and high-quality grains (4, 5). Colombia’s main cocoa-producing regions include Santander, Antioquia, Arauca, Tolima, and Huila. In 2020, cocoa production generated 165,000 direct and indirect jobs in Colombia, making it one of the primary sources of income for over 52,000 Colombian families. Furthermore, cocoa has been established as an alternative to peace efforts in post-conflict territories and as a substitute for illicit crops (4). Cocoa is considered to play an essential role in Colombia’s social and economic development.

However, climate change can affect cocoa production, primarily because this crop is sensitive to abiotic stresses such as drought and salinity (1). This crop requires temperatures between 18°C and 32°C, 1,500 and 2,500 mm of rainfall, and constant humidity for optimal growth (5, 6). The 2024 report by United Nations indicates an increase in the frequency of daily heatwaves and drought months in recent decades, which may lead to a decline in agricultural production and heightened food insecurity (7). Additionally, the 2024 report from the Colombian Institute of Hydrology, Meteorology, and Environmental Studies (IDEAM) predicts variable drought events across the country’s cocoa-producing regions (8). For departments such as Antioquia and Huila, an increase in precipitation is expected; however, in Arauca, Santander, and specific areas of Tolima, a decrease in rainfall is anticipated, resulting in longer drought seasons (8). Considering that cacao seedlings and juveniles may be more susceptible to drought due to their shallow rooting depth, which ranges from 0.4 to 0.8 meters, water absorption may be severely limited during extended drought periods (1, 9). This, in turn, adversely affects nutrient uptake, water relations, and gas exchange. As a result, these conditions may lead to alterations in the crop’s growth, biomass, and various biochemical characteristics, ultimately impacting cocoa productivity and quality (9)

Different strategies have been proposed given the effects of drought on cocoa production. One potential strategy is using endophytic fungi (10), microorganisms that live inside the plant without causing apparent damage (11). On the contrary, endophytic fungi can confer several benefits to the plant, including protection against herbivory and plant pathogens, improved growth, protection, and tolerance to abiotic stress (12, 13). Endophytes can stimulate plant responses, such as producing reactive oxygen species (ROS) enzymes and increasing phytohormone levels, which promote root and shoot growth (14, 15). Consequently, fungal endophytes could represent an important alternative to enhance the overall fitness of the plant under drought conditions (10). However, it is essential to note that only 1% of environmental microorganisms have been identified, underscoring the need for a deeper understanding of the diversity of microorganisms, such as endophytes (16). These microorganisms represent a group with great potential, and further exploration could uncover new possibilities for their application in agriculture. Studies on microbial diversity in soil and endophytes have shown their importance. For instance, research by Erktan et al. suggests that a high diversity of these microorganisms increases the likelihood of discovering species with various potential agricultural applications, such as nutrient assimilation, biocontrol, growth promotion, protection against abiotic stress, and water content regulation, among others (17).

Xerophytic angiosperms, exemplified by cacti, have been identified as hosts to endophytes. These endophytes have been associated with mitigating abiotic stresses, such as salinity and drought, and fostering the growth of host plants (18). According to studies on drought tolerance in tomato and cucumber, such as the one developed by Miranda et al. tomato plants inoculated with endophytes from *Eragrostis cilianensis* showed optimal tolerance to drought stress (19). These plants exhibited lower levels of ROS enzymes, higher stomatal conductance, and increased root biomass, among other beneficial effects, compared to the control group. Therefore, the endophytes residing within cacti, such as *Stenocereus* spp. emerge as a potential strategy for inducing drought tolerance in crops.

To address the pressing challenges posed by climate change on cocoa production, this study aimed to isolate endophytic fungi from the roots of *Stenocereus* spp. plants located in two arid zones of Colombia, Tatacoa (Huila) and Taganga (Magdalena), and to assess the diversity of these fungi. This allowed us to evaluate their potential role in enhancing drought tolerance in *T. cacao*. It also contributed to better understanding fungal diversity and its potential applications in sustainable agricultural practices.

## Methods and materials

### Study site and sampling

Endophytes were sampled in July 2023 in two localities in Colombia: The Tatacoa Desert of the municipality of Villavieja, Huila department, and Taganga, Magdalena department (see all geographical coordinates in Supp. Table 1). At each location, roots were collected from 12 mature cactus *Stenocereus* spp. specimens at depths ranging from 1to 20 cm, with a minimum separation distance of 7 m between sampling points. Following the ensuing collection, specimens were hermetically sealed and stored at 10 °C for 72 h (20). Subsequent processing for endophytic fungi isolation occurred at the Mycology and Phytopathology Laboratory (LAMFU) of the Universidad de los Andes (Bogota, Colombia).

**Table 1.** Diversity metrics for two localities, Tatacoa and Taganga. Confidence intervals (IC Inf, IC Sup) indicate the range of uncertainty for the Shannon index.

| Locality | Species richness | Shannon |  | Simpson | Fisher | Simpson's inverse |
| --- | --- | --- | --- | --- | --- | --- |
|  |  | IC Inf | IC Sup |  |  |  |
| Tatacoa | 39 | 2.871 | 3,50 | 0.933 | 18.194 | 14.871 |
| Taganga | 31 | 2.570 | 3.27 | 0.912 | 14.014 | 11.321 |

Soil samples (two samples per locality) were collected at a depth of 5 cm from each locality, Tatacoa and Taganga. Laboratory analyses were performed to determine salinity and pH (18).

### Isolation of fungal endophytes

A systematic procedure was implemented to isolate endophytic fungi from the roots of each plant, involving a thorough wash with tap water to remove any residues of plant material and soil. Subsequently, a disinfection process was done following the protocol Silva-Hughes et al. (20). This involved immersing each sample in 70% ethanol for one minute, followed by a three-minute wash with 2% hypochlorite and a subsequent rinse with sterile distilled water (SDW). Noteworthy modifications to the protocol included an additional wash with 70% ethanol after hypochlorite treatment, followed by a final rinse with SDW. The roots were meticulously dried using sterile absorbent paper after the washing procedure.

To assess the efficacy of the procedure, disinfected roots were imprinted on Petri plates with Potato Dextrose Agar supplemented with chloramphenicol (500 mg/l) (PDA-C) (21). Subsequently, 0.5 cm segments were excised from the disinfected roots, and five fragments were placed on Petri plates containing PDA-C. The plates were incubated for up to 7 days at 30 °C. This entire process was replicated in triplicate for each plant (20). An axenic culture of each fungus from the root was then established in PDA-C and further incubated at 25 °C (20).

### Identification and diversity of fungal endophytes

The fungal endophyte isolates were grouped into morphotypes, or genera, when possible, based on macroscopic description and molecular identification. For the morphological description, the colony of each isolate was described based on its macroscopic characteristics, including surface texture, color (color code: https://color.bio/color-picker), aerial mycelium, and structure production. A detailed microscopic description was made only for the morphotypes identified by molecular methods, following the dichotomic key of Cepero et al. (22), Carmichael (23), and Seifert and Gams (24). All pure isolates were preserved in vials with sterile water.

For molecular identification, fungal isolates were grown on PDA for 5 to 8 days at 25 °C. DNA was extracted using the modified SDS method of Natarajan et al. (25): the lysis was performed in an Eppendorf tube with beads and 500 µl of lysis buffer (100 mM Tris, pH 8.0, 50 mM EDTA, and 1% SDS). One gram of the colony was added, and then subjected to a bead beater (Mini-Beadbeater-16) for 40 sec. For purification, 275 µL of ammonium acetate (7 M, pH 7.0) and 500 µL of chloroform were added, with an incubation of 40 min at 68 °C between the two components. Lastly, DNA precipitation was achieved by adding 1 mL of isopropanol and incubating for 24 h, followed by a final wash with 1 mL of 70% ethanol. The extracted DNA was eluted with 50 µL of Milli-Q water and stored at -20 °C.

All isolates were identified using the internal transcribed spacer region of the rRNA (ITS) with the universal primers ITS1 and ITS4 (26). For the *Fusarium* genus, we also used the translation elongation factor 1-alpha (*Tef-1a*) (27). Single-band PCR products were sequenced using an ABI 3500XL DNA analyzer in the Sequencing Core Facility (Gencore) at the Universidad de Los Andes (Bogotá, Colombia). Sequence assembly and editing were performed using Geneious software v. 2022.2.2 (https://www.geneious.com). Preliminary identification of the isolates to the genus level was performed by analyzing gene sequences, using the BLASTn algorithm with default parameters and choosing the type organisms (28).

Diversity metrics were calculated using the *vegan* package of R Software version 4.4.3 (29) (30). Alpha diversity was estimated with Shannon Fisher α, Simpson indexes, and the inverse of the Simpson index (31, 32). Furthermore, the bootstrap method was employed to estimate the uncertainty of the Shannon index (20).

### In-vitro drought tolerance evaluation

For the in vitro drought tolerance assay, 22 purified morphotypes were selected based on PDA and propagule production growth rates. The strains were incubated on PDA medium at 25 °C. Two plugs of the actively growing endophytic fungi were used to inoculate potato dextrose broth (PDB) supplemented with polyethylene glycol (PEG-6000) at a concentration of 20% w/mL, which generates an osmotic pressure of -0.6 MPa (33). Additionally, as a control, two plugs of the actively growing endophytic fungi were used to inoculate a PDB medium without PEG-6000. Subsequently, all the cultures were incubated in a shaker for 7 days at 28 °C and 160 rpm (33). The experiment was performed in triplicate for each morphotype. Finally, the percentage of biomass loss (equation 1) was calculated to determine the drought tolerance of the morphotypes. Fungal strains exhibiting less than 20% biomass loss were selected for in vivo assays:

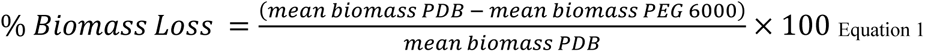

### Plant growth conditions, inoculation, and drought stress assay

Based on the previous in vitro drought tolerance evaluation, five fungi were selected due to their higher biomass production in PDB-PEG6000 and a less than 20% biomass loss. The five chosen fungi were used to evaluate drought tolerance in 5-month-old juvenile *T. cacao* plants under greenhouse conditions at Pitalito-Huila (1°52’07.3”N 76°03’50.8” W) during June-July of 2024. Abiotic conditions: air temperature, soil water content (SWC), relative humidity (RH), and photosynthetically active radiation (PAR) were monitored in the greenhouse using the corresponding sensors attached to a HOBO datalogger (model H21, Onset). The experimental design used for this study was randomized. The 5-month-old cocoa juvenile plants, with an average height of 37.4 cm and approximately 12 leaves, were obtained from Maloka Cacao Nursery located in Rivera-Huila, planted in black polyethylene bags (15 x 30 cm and 63 mm caliber) with 2 kg of non-sterile substrate (6 soil:1 husk:1 ash) (34).

A total of 180 juvenile plants were used, with 90 assigned to the drought treatment (15 per five fungal inoculation treatments, and 15 for a control with SDW supplemented with 1% Tween 80) and 90 to the non-drought control. A culture grown on PDA and oatmeal medium at 25 °C for 15 days of each morphotype was used to prepare the inoculum suspension. The spore suspension was adjusted to 1 × 10^6^ conidia/mL in SDW supplemented with 1% Tween 80 (35, 36). The substrate was irrigated with 125 mL of fungal inoculum per 2 kg substrate for each inoculation treatment.

Over six days, plants were acclimated in the greenhouse and irrigated with tap water at 7:00 a.m. On day 6, the plants were inoculated with the different fungal strains, and three days post-inoculation (day 9), the drought stress assay commenced. A total of 15 plants per fungal morphotype and non-inoculated plants were exposed to drought stress for 13 days (days 10 to 22), during which they began to exhibit apparent symptoms and signs of drought-related damage. After 13 days (day 23), the plants were re-irrigated with tap water under normal conditions for two weeks. The plants in the non-drought control group continued to be irrigated with tap water at 07:00 h, three times a week.

### Physiological parameters

Leaf water potential (Ψ_L_) and stomatal conductance (Ks) were measured on mature leaves located below the third node of juvenile plant samples, between 11:00 and 14:00 h, at three time points during the experiment: (i) three plants from each treatment, before the drought cycle began (5 days before inoculation); (ii) five plants from each treatment, 13 days after the drought stress cycle started, and (iii) three plants from each treatment, two weeks after the plants were re-irrigated. Ψ_L_ was measured following the protocol outlined by Dos Santos et al. using a pressure chamber (model 1505D-EXP, PMS Instruments) (37). Ks was measured with a leaf porometer (SC-1, Decagon Devices) (38).

### Measurement of morphological and growth parameters

Plant height, number of leaves, specific leaf area (SLA), leaf area, and biomass were measured at three time points as described previously. Plant height, number of leaves, and leaf area were measured for each treatment according to the protocol of (36, 39).

Biomass was determined according to the modified protocol of Dos Santos et al. in which the roots, stems, and leaves of the plants of each treatment were stored, separated in paper bags, and oven dried at 75 °C (Memmert UM300) for 72 h until constant dry weight (g) (37). Furthermore, the relative growth rate (RGR) was determined according to Equation 2 (40), where S_1_ and S_2_ correspond to initial and final biomasses, respectively, at times 1 (t_1_) and 2 (t_2_).

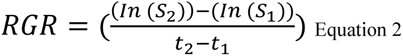

According to Roderick et al. specific leaf area (SLA, cm^2^/g) was determined by collecting the third fully developed leaf with a similar leaf index color (41). Subsequently, the leaf samples were wrapped in aluminium foil and stored until processed (42). ImageJ software measured leaf area (cm^²^) (43). Leaves were then oven-dried at 70 °C (Memmert UM300) for 72 h until constant dry weight (42).

### Measurement of proline content

The activity of free proline (µg/mg) was evaluated at the beginning of the experiment (day 0) and after 13 days of exposure to both drought and non-drought control conditions (day 23). A calibration curve was performed using known proline content to ensure accurate quantification. The proline content was determined from healthy leaves of *T. cacao* juvenile plants following the methods described by Bates et al. (44) and Dos Santos et al. (37). Healthy leaves of *T. cacao* ICS95 were stored at –80 °C until extraction. The leaf tissue was lyophilized for 48 h, after which it was ground in liquid nitrogen. Approximately 35 mg of plant material was mixed with 1 mL of 3% sulfosalicylic acid, followed by agitation in a shaker at 240 rpm for 30 min and centrifugation at 14,000 x g for seven min. Subsequently, 0.2 mL of the crude extract was transferred to an Eppendorf tube along with 0.2 mL of acid ninhydrin (1.25 g ninhydrin, 30 mL glacial acetic acid, and 20 mL 6 M phosphoric acid) and 0.2 mL of glacial acetic acid (37, 44). The mixture was vortexed for 15 sec. The Eppendorf tubes were incubated in a water bath at 85 °C for 120 min. Immediately afterward, the reaction was stopped by placing the tubes in an ice bath for 20 min. Finally, 0.4 mL of toluene were added to the tubes and vortexed for 15 sec (Dos Santos et al., 2020). The samples were left at room temperature to allow phase separation. Absorbance readings were taken at 520 nm using a spectrophotometer (Elisa SpectraMax M4), in triplicate, with toluene as the blank (37).

### Statistical analyses

We first assessed normality and homogeneity of variances. Depending on these results, a t-test or a Wilcoxon test was conducted to evaluate differences between juvenile plants under drought and non-drought conditions. Additionally, ANOVA or Kruskal–Wallis tests, followed by appropriate post hoc analyses, were performed to compare the effects of different treatments on juveniles inoculated with fungal endophytes and on non-inoculated controls under each condition. All analyses were conducted in R software version 4.4.3 (30).

## Results

### Soil analyses

Soil samples collected from the Tatacoa (n= 2) showed electrical conductivity (EC) values of 30 and 69 ppm, with pH values of 9.6 to 9.9. In contrast, samples of Taganga locality (n= 2) exhibited EC values of 118 and 119 ppm and pH values of 8.8 and 8.9.

### Isolation of fungal endophytes and morphotype characterization

A total of 308 fungal endophytes (n=170 for Tatacoa and n=138 for Taganga) were isolated. Forty-nine morphotypes were identified in both localities; 21 were exclusive to Tatacoa and 14 to Taganga. All morphotypes were identified based on their phenotypic characters (Supp. Table 2). Molecular identification also revealed the presence of 49 potential different taxa (Supp. Table 2). Of these, 31 were found in both localities, while 12 were exclusive to Tatacoa and six to Tatacoa. Among the genera identified based on morphological and molecular characteristics in both localities, prominent taxa included: *Alternaria* sp., *Aspergillus* spp., *Curvularia* spp., *Ectophoma* sp., *Epicoccum* sp., *Fusarium* spp., *Lasiodiplodia* sp., *Macrophomina* sp., *Moniliophthora* sp., *Monosporascus* sp., *Phoma* sp., *Rhizopus* sp., and *Trichoderma* spp. Furthermore, the genera *Acrophialophora* sp., *Latorua* sp., *Myrmaecium* sp., *Polyporus* sp., and *Rhodotorula* sp. were uniquely found in the Taganga locality. In contrast, *Acrocalymma* sp., *Clavulina* sp., *Coprinellus* sp., *Deniquelata* sp., *Didymella spp*., *Diaporthe* spp., *Didymocrea* sp., *Oblongocollomyces* sp., *Paraboeremia* sp., *Phoma* sp., and *Purpureocillium* sp. were exclusively identified in the Tatacoa locality.

**Table 2.** Mean leaf water potential (Ψ_L_, MPa) and stomatal conductance (Ks, mmol/m^2^s) with standard deviation for the treatments at the beginning (day 0), after 13 d under drought conditions (day 23), and after re-watering the juvenile plants for two weeks (day 34). Statistical differences between treatments are represented by 0 ‘***’ 0.001 ‘**’ 0.01 ‘*’ 0.05 ‘.’ 0.1 ‘’ 1.

| Treatment | Day 0 |  | Day 23 |  | Day 34 |  |
| --- | --- | --- | --- | --- | --- | --- |
| | $\Psi_L$ | Ks | $\Psi_L$ | Ks | $\Psi_L$ | Ks |
| *Ectophoma* sp. | -0.92<br>±0.11 | 262.97 ±45.65 | -5.6 ± 0.4 | 40.73 ± 11.42 | -5.73 ± 0.45 | 40.73 ± 11.42 |
| *Acrophialophora*<br>sp.1 | -<br>1.02 ± 0.25 | 250.7 ± 64.76 | -<br>4.82 ± 1.06 | 39 ± 17.91 | -5.74 ± 0.36 | 39 ± 17.91 |
| *Fusarium* sp. | -<br>0.96 ± 0.18 | 189.9 ± 32.22 | -<br>4.88 ± 1.28 | 39.77 ± 7.65 | -5.22 ± 0.41 | 39.77 ± 7.65 |
| *Phoma* sp. | -0.96 ± 0.2 | 161.53 ± 10.22 | -<br>4.55 ± 1.03 | 31.32 ± 6.85 | -<br>3.98 ± 0.74* | 32.17 ± 8.13 |
| *Didymocrea* sp. | -<br>1.03 ± 0.15 | 217.6 ± 66.02 | -<br>5.36 ± 1.07 | 44.87 ± 15.7<br>2 | -5.37 ± 1.37 | 44.87 ± 15.72 |
| Non-inoculated | -<br>0.97 ± 0.17 | 213.37 ± 58.93 | -<br>5.92 ± 0.23 | 39.2 ± 11.86 | -5.99 ± 0.24 | 39.2 ± 11.86 |

The ecological analyses showed differences between the two localities. Tatacoa exhibited higher diversity compared to Taganga (Table 1). Furthermore, Tatacoa had higher values across all diversity indices: Shannon index (2.871); Simpson index (3.50), indicating a higher level of evenness; and Fisher’s alpha (18.194), reflecting a richer community. Cacti roots in both localities generally exhibited high levels of fungal diversity.

### In-vitro drought tolerance evaluation

Twenty-two taxa were selected for an *in vitro* drought assay based on growth rates and the production of propagules (Fig. 1). Of the 22 fungal strains tested, 11 experienced a biomass loss of less than 20% (red line in Fig. 1A). Nine of these 11 strains showed higher biomass under drought conditions compared to normal conditions. Subsequently, the biomass of the eleven morphotypes under drought conditions was evaluated (Fig. 1B). Based on these results, five strains: *Acrophialophora* sp., *Didymocrea* sp., *Ectophoma* sp., *Fusarium* sp.1, and *Phoma* sp. were selected for the greenhouse drought assay because they showed the highest biomass under drought conditions and experienced less than 20% biomass loss (Fig. 1).

**Figure 1.**
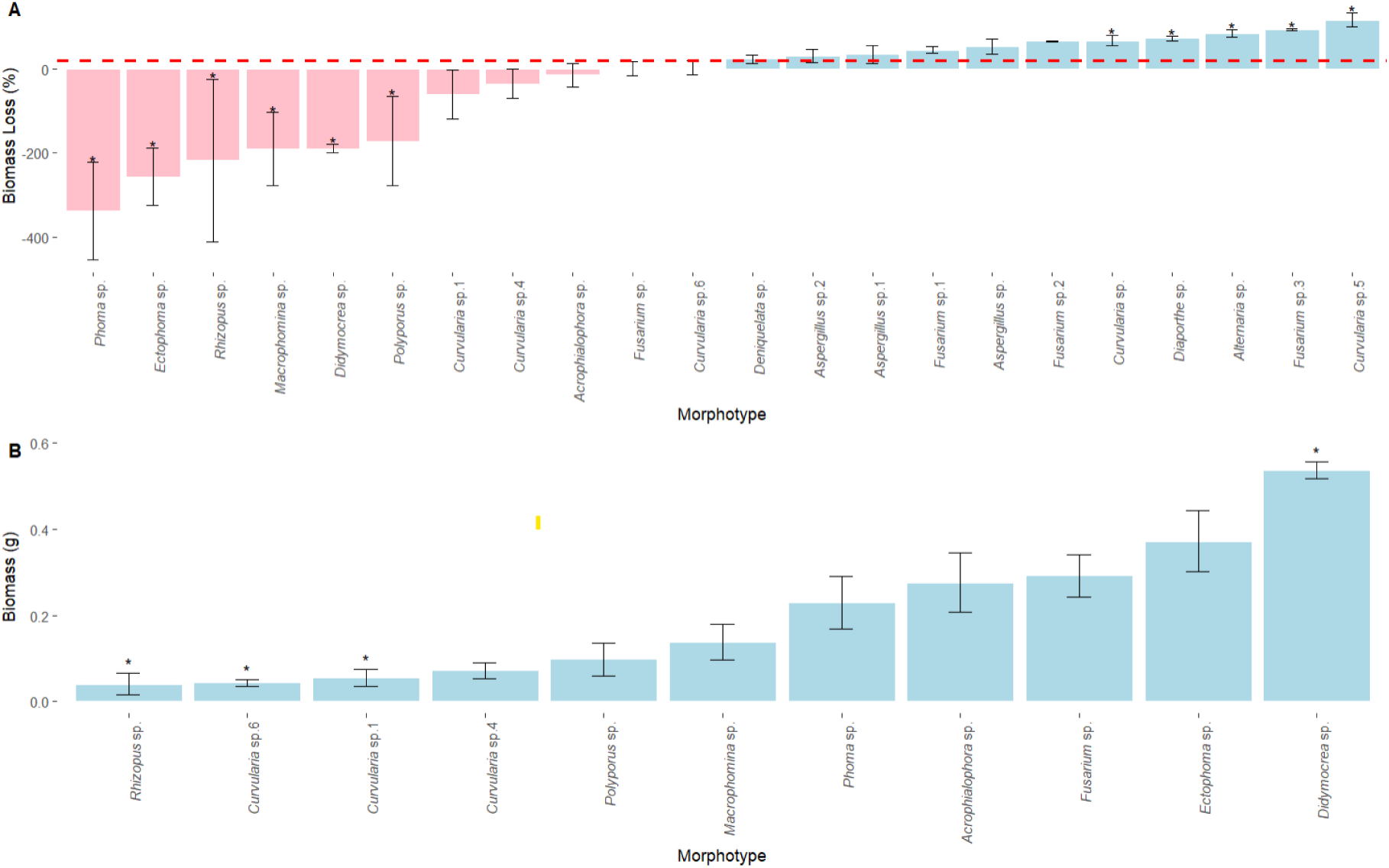
Biomass analysis of fungal endophytes under normal conditions (PDB) and drought conditions (PDB+PEG 6000). A. Percentage of biomass loss of fungal endophytes. The horizontal red line shows a reduction in biomass of 20%. B. Biomass of the 11 morphotypes that showed less than 20% biomass loss under drought conditions. Bars show mean ± SE. Asterisks denote statistical significance among morphotypes based on Kruskal– Wallis followed by Dunn’s test (p < 0.05).

### Drought stress assays

#### Soil Water Content

Before the drought stress treatment, plants were irrigated and started with similar volumetric water (0.1121 - 0.1561 m³/m³) (day 0). Moreover, after 10 days of drought (day 19), the soil water content (SWC) of juvenile plants exposed to drought remained constant for the last 4 d (less than 0.05 m³/m³) of the drought stress assay (Fig. 2). After re-watering for 11 days, the SWC of juvenile plants under drought conditions increased, reaching values of 0.1252-0.1275 m³/m³ (Fig. 2).

**Figure 2.**
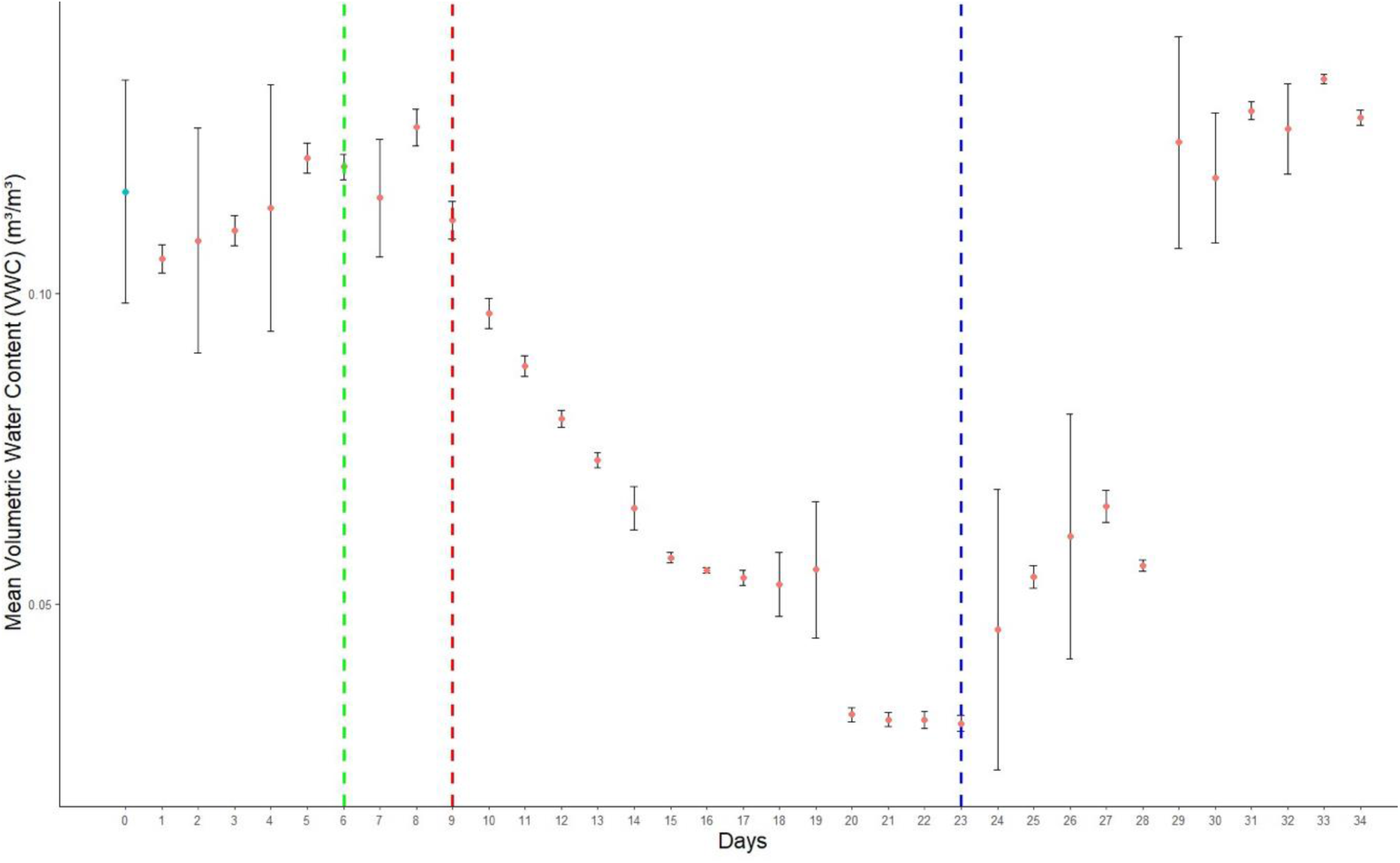
Mean soil water content (SWC, m³/m³) over time, measured daily, with error bars indicating standard deviation. Green vertical line on day 6 indicates inoculation of the soil with the fungal endophytes. Red vertical line on day 9 indicates the onset of drought stress. Blue vertical line at day 23 indicates the end of drought stress and the beginning of the re-irrigation.

#### Temperature, relative humidity, and photosynthetic active radiation

Within the greenhouse, the average temperature was 26.5 °C, the relative humidity (RH) was maintained at 72.90%, and the photosynthetically active radiation (PAR) level averaged 147.042 µmol/m²/s (see all data in Supp. Table 3). However, temperature and RH varied significantly during the periods when leaf water potential (ΨL) and stem hydraulic conductivity (Ks) were assessed (0, 23, and 34 days). Temperature and RH were higher after 11-13 days of drought exposure (days 21, 22, and 23) and following the two-week rehydration period (day 34) compared to day 0 of the experiment. (Temperature: p-value < 0.001; RH: p-value < 0.001; α = 0.05). Finally, significant differences in PAR were also detected when we measured it at day 34 (p = 0.009; α = 0.05), following the rehydration period (Supp. Table 3), compared to day 0.

#### Physiological response: Leaf water potential (Ψ_L_) and stomatal conductance (Ks)

Leaf water potential (Ψ_L_) and stomatal conductance (Ks) were comparable across all treatments at the beginning of the experiment (day 0), showing no significant differences (Ψ_L_: p-value = 0.845, α = 0.05; Ks: p-value = 0.217, α = 0.05) (Fig. 3). By day 23, juvenile plants subjected to drought stress exhibited a significant reduction in both Ψ_L_ and Ks compared to those under non-drought conditions (p-value < 0.001, α = 0.05) (Table 2 and Fig. 3). Moreover, no significant differences were observed between juvenile plants inoculated with fungal endophytes and non-inoculated plants under drought conditions after 13 d, with p-values of 0.251 for Ψ_L_ and 0.153 for Ks (α = 0.05) (Fig. 3).

**Figure 3.**
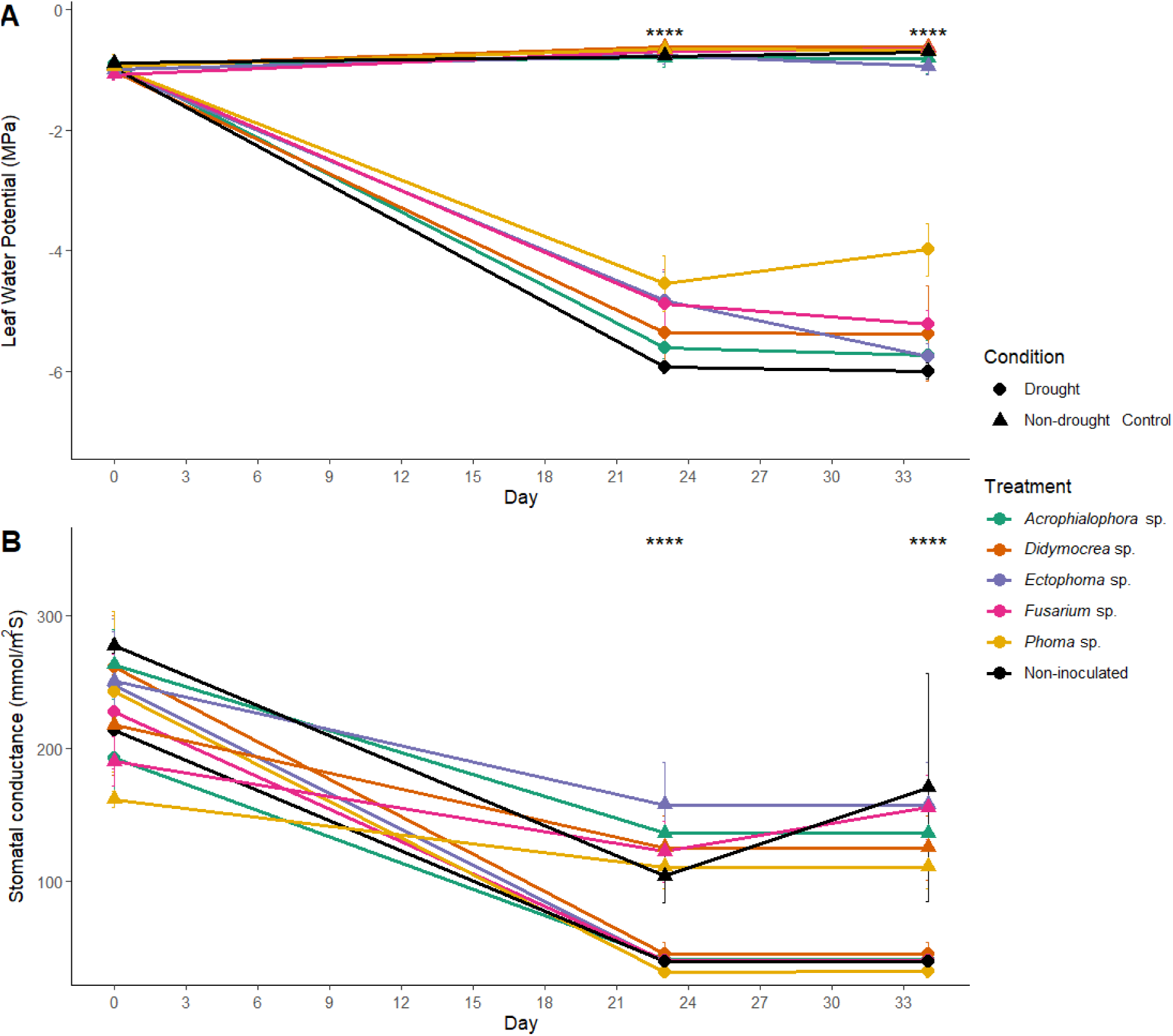
Physiological responses of *Theobroma cacao* ICS95 juvenile plants inoculated with fungal endophytes and non-inoculated plants subjected to drought conditions and non-drought control. Day 0 represents non-inoculated plants under normal conditions. Day 23 corresponds to plants inoculated and subjected to 13 days of drought stress. Day 34 shows plants after two weeks of rewatering. (A) Mean leaf water potential (Ψ_L_) with error bars. (B) Mean stomatal conductance (Ks) of juvenile plants with error bars. The black lines indicate the mean Ψ_L_ or Ks of the treatment under non-inoculated control conditions. Asterisks denote statistical significance.

Under non-drought control conditions, Ks decreased after 23 and 34 days of the experiment compared to day 0 (p-value = 0.0025, α = 0.05). Juvenile plants inoculated with the five fungal endophytes experienced a decline in Ks on day 23, which stabilized by day 34. Meanwhile, non-inoculated juveniles showed a decrease in Ks until day 23 after 13 days of drought, followed by an increase on day 34 (p-value = 0.0927, α = 0.05) (Fig. 3).

Finally, on day 34, two weeks after rewatering, most juvenile plants failed to recover from drought stress. An exception was observed in treatment with the fungus *Phoma* sp., which exhibited a significantly higher Ψ_L_ (Fig. 3) and a foliar recovery of 57.14%.

#### Growth parameters, relative growth rate, and specific leaf area

After the drought experiment period, during the recovery and rewatering period, the relative growth rate (RGR) in non-drought control conditions decreased in juveniles inoculated with fungal endophytes, except for those inoculated with *Phoma* sp., which exhibited a trend toward a higher RGR (Fig. 4). However, this difference was not statistically significant (p-value = 0.431, α = 0.05). Juveniles, both inoculated and non-inoculated with fungal endophytes, that were exposed to drought conditions demonstrated a significant decrease in RGR compared to those under normal conditions, as indicated by the Kruskal-Wallis test (p-value = 0.009359, α = 0.05). Furthermore, juveniles inoculated with *Ectophoma* sp. and *Fusarium* sp.1 exhibited a lesser decrease in RGR compared to non-inoculated juveniles under drought conditions. However, this difference was also not significant (p-value = 0.998, α = 0.05) (Fig. 4).

**Figure 4.**
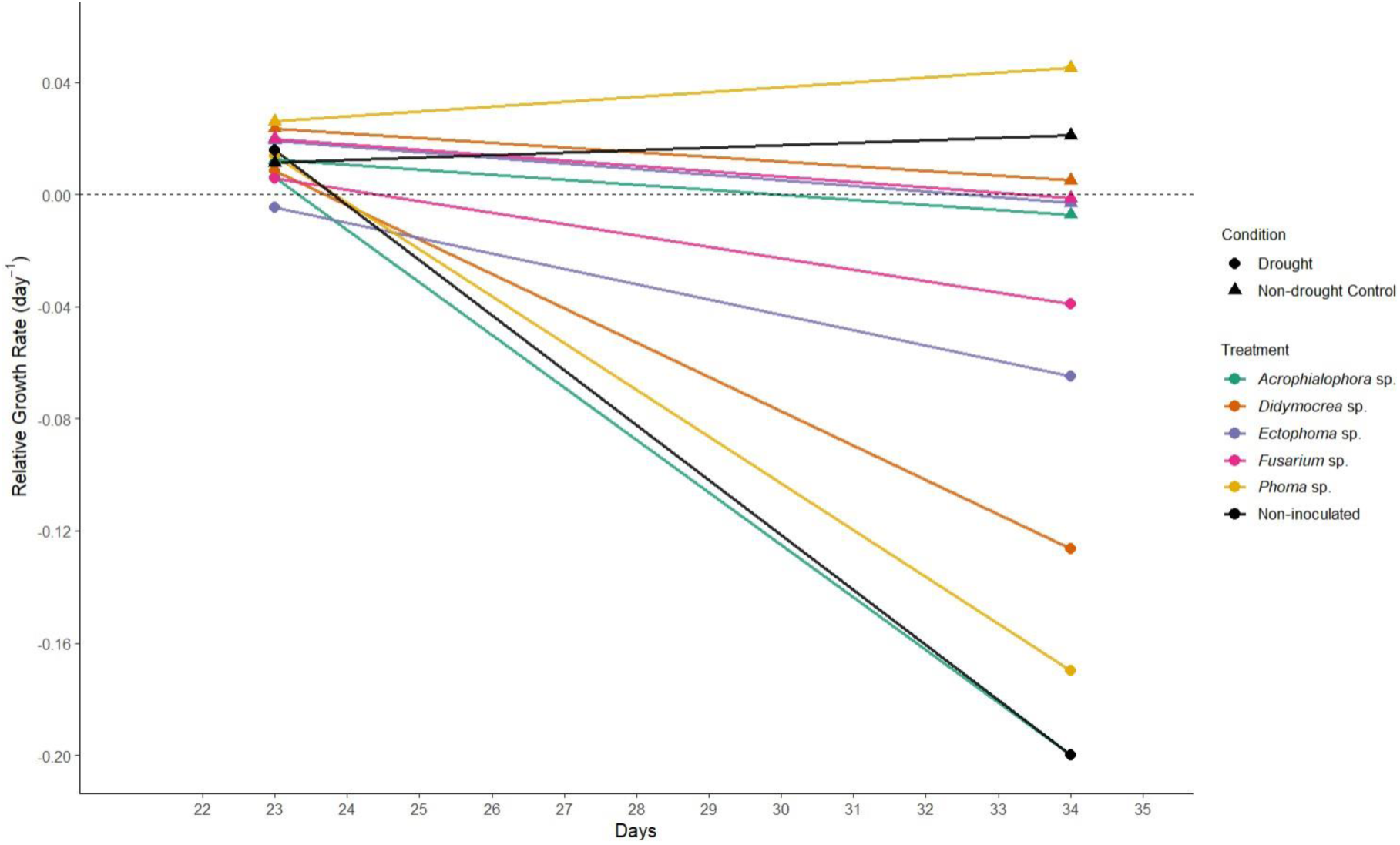
Relative Growth Rate (day^-1^) of juvenile plants of *Theobroma cacao* ICS95 inoculated and non-inoculated with the fungal endophyte *of Stenocereus* spp. during the rewatering period. Growth rates are compared under non-drought control and drought.

Regarding plant height, no significant differences were observed among treatments throughout the experiment (p-value = 0.5457, α = 0.05). However, significant variations (p-value = 0.0012, α = 0.05) were noted in the number of leaves between juvenile *T. cacao* plants subjected to drought stress and those in normal conditions. Specifically, drought-stressed plants exhibited a significant reduction in leaf count and specific leaf area (SLA) (Supp. Table 2) (p-value < 0.001, α = 0.05). In contrast, among plants under non-drought conditions, those inoculated with the fungus *Fusarium* sp. 1 showed a significant increase in SLA (p-value < 0.001, α = 0.05) at both 23 and 34 d compared to non-inoculated plants and baseline measurements (Supp. Table 4).

Lastly, juveniles under drought conditions showed a significant decrease in the area of new leaves compared to the control group. In contrast, juveniles inoculated with *Phoma* sp. showed a slight (non-significant) increase in leaf area compared to the control at day 23 (Supp. Fig. 1) and Fusarium at day 34 (Supp. Fig. 2).

#### Proline content

At day 0, no significant differences in proline content were observed between treatments. However, after 13 days of drought stress (day 23), a significant increase in leaf proline content was observed compared to those under non-drought control conditions (p-value < 0.001, α = 0.05). This increase in proline content, indicative of drought tolerance, was particularly pronounced in plants inoculated with *Acrophialophora* sp., *Fusarium* sp. 1, and in the non-inoculated group (Fig. 5). In contrast, treatments with *Phoma* sp., *Ectophoma* sp., and *Didymocrea* sp. exhibited a significantly smaller increase in proline levels compared to the non-inoculated group (F-statistic = 8.274, p-value <0.001, α = 0.05). Interestingly, proline accumulation was also significantly lower in plants inoculated with *Phoma* sp. and similar to not subjected to drought.

**Figure 5.**
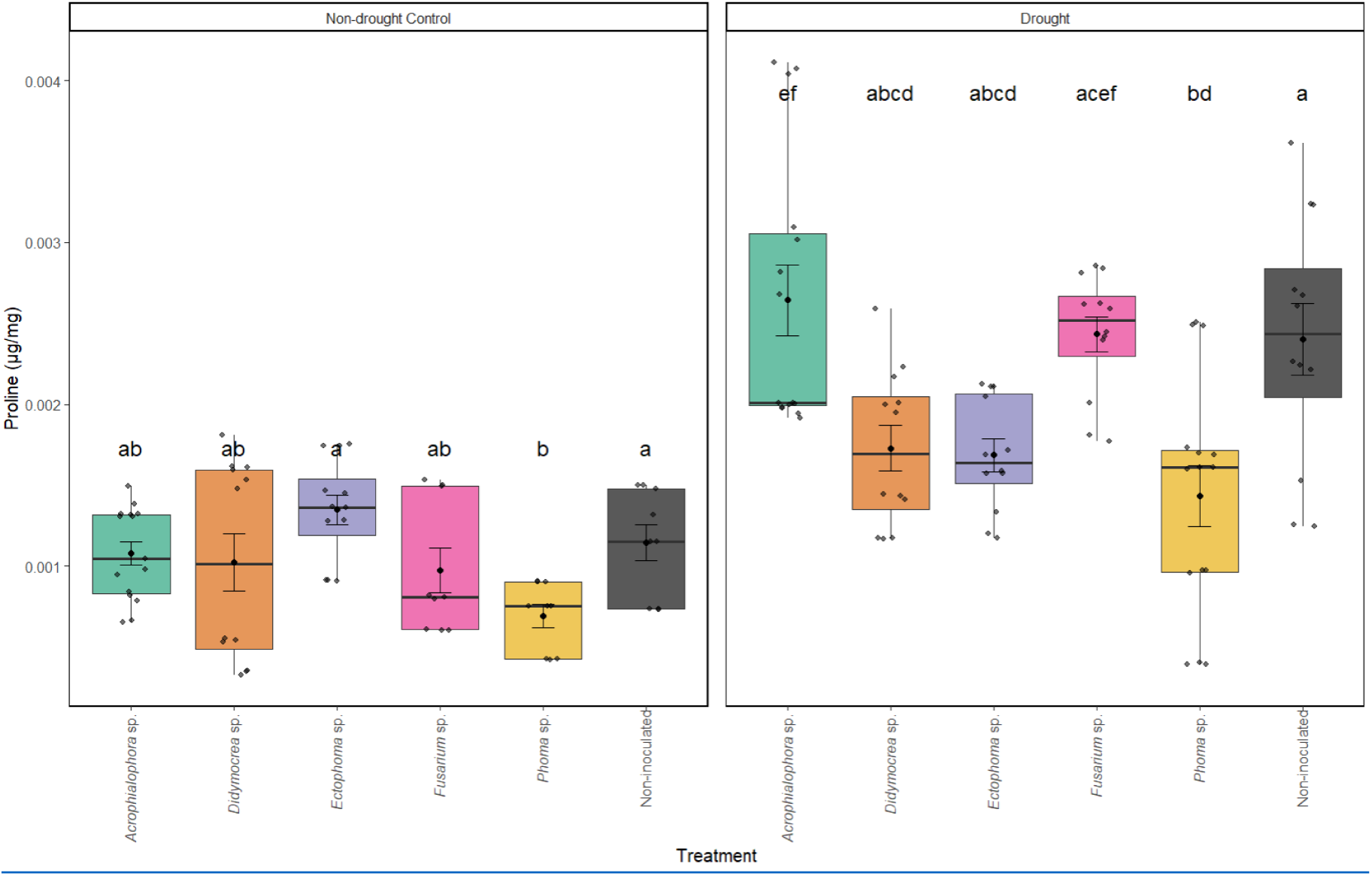
Free proline content (μg/mg) in leaves of juvenile plants of *Theobroma cacao* ICS95 after 13 d of non-drought control (leaf) and drought (right). Letters denote statistical significance using the Kruskal–Wallis test followed by Dunn’s post hoc test for the non-drought control, and ANOVA followed by Tukey’s test for the Drought (p < 0.05).

## Discussion

This study represents the first report on isolating and evaluating fungal endophytes from *Stenocereus* spp. in relation to their effects on *T. cacao*. Our findings highlight the potential applications of these microorganisms to enhance agricultural resilience and sustainability. The endophytes isolated from *Stenocereus* spp. demonstrated the capacity to promote plant growth and improve drought tolerance in juvenile plants of *T. cacao* ICS95. Some fungal genera contributed to plant recovery following drought stress. According to Yarzábal Rodríguez et al. (45), extremotolerant fungi may exhibit an enhanced ability to confer drought tolerance in crops. Nonetheless, care must be taken with extremophilic fungi that can adapt to higher temperatures and show undesirable pathogenicity (45). Additionally, specific genera isolated in this study have previously been reported as plant pathogens. Therefore, the eventual deployment of these strains in crops should be complemented by a pathogenic host range study.

The cacti roots represent a promising niche for discovering diverse fungi that exhibit various phenotypes that enhance plant resilience. Some genera identified in this study, including *Alternaria* spp., *Aspergillus* spp., and *Fusarium* spp., are among the most frequently isolated fungi from plants in the Cactaceae family (46, 47). Additionally, records of *Diaporthe* spp. and *Rhodotorula* spp. have been noted in the study of Ferreira-Silva et al. on endophytic fungi in *Melocactus ernestii* (48). Our study is the first to report *Polyporus* spp. as an endophyte in cacti, although previously identified as capable of growing in xerophytic environments (49).

We found high levels of fungal diversity in the roots of cacti in both localities. Diversity of microbial communities may be shaped by several soil characteristics, particularly aridity (18) and pH (50). Similarly, Loro et al. suggest that the diversity of endophytic fungi isolated from arid environments can be influenced by factors such as the type of isolation method used and the sample size (47). In this study, the main factors that may influence fungal diversity levels are the isolation protocol employed and the pH levels of the sampled environments. The reported electrical conductivities of the Tatacoa and Taganga classify them as non-saline, suggesting that microbial respiration is not inhibited. However, the pH values were higher than eight in both locations, indicating that the soils are alkaline and characteristic of arid or semi-arid environments (50). Microorganisms growing in soils with high pH values can be considered either alkaliphilic or alkalitolerant (51). It is important to note that the diversity observed at each location may be underestimated, as it was determined solely based on culturable endophytic fungi isolated using a single growth medium (PDA). To enhance the understanding of fungal diversity present at each locality, complementary metagenomic studies are recommended, alongside the use of diverse culture media and techniques that facilitate or optimize the isolation of xerophilic and alkaliphilic fungi (52).

Several endophytes, including *Acrophialophora* sp., *Didymocrea* sp., *Fusarium* sp., and *Phoma* sp., showed the ability to grow in vitro under intense drought stress. Interestingly, these genera have been previously reported as microorganisms with the potential to induce drought tolerance in various crops (53–56). Additionally, it has been reported that *Acrophialophora* sp., *Didymocrea* sp., *Phoma* sp., and *Fusarium* sp. may play a significant role in protection against plant pathogens, as well as exhibiting potential activity as growth promoters (53, 54, 56, 57).

Three approaches were employed to assess endophytic fungi’s role in inducing drought tolerance in *T. cacao*: physiological responses, relative growth rates, and biochemical responses to water stress. In the first approach, it was observed that juveniles of *T. cacao* ICS95 exposed to drought conditions reached Ψ_L_ values below -5.0 MPa, maintaining them above the fatal failure. Seleiman et al. reported that values below -6.0 MPa can result in fatal hydraulic failures in plants (58). On the other hand, it is worth noting that the Ψ_L_ values obtained in this study are lower than those reported by Osorio-Zambrano et al. for *T. cacao*, who found values of -3.0 and -3.5 MPa in juveniles under drought conditions (1). This discrepancy may be attributed to the fact that measurements in the Osorio-Zambrano et al. study were taken during predawn hours, corresponding to more favorable conditions (1). In contrast, our measurements were recorded between 11:00 and 13:00 h, which is associated with more stressful plant conditions. Moreover, the experiment conducted by Osorio-Zambrano et al (1) lasted for 26 days, in contrast to the 13 days of water stress applied in this study, after which the plants began to experience decay and wilting. This difference in results could be attributed to the microclimatic conditions under which both experiments were conducted and the amount of substrate used. In this study, the substrate for juveniles was 2 kg, while Osorio-Zambrano et al. used 5 kg (1). Less substrate means less water retention, reducing plant water availability (59).

*Theobroma cacao* ICS95 juveniles subjected to drought conditions significantly decreased Ψ_L_ compared to those under optimal conditions. This phenomenon is attributed to the reduction of available water in the soil, which causes an increase in the concentration of solutes within the plant (60, 61). As a defense mechanism, individuals close their stomata to minimize water loss, which explains the significant decrease in stomatal conductance (Ks) compared to the control group (60, 62). However, stomatal closure reduces photosynthetic rate, which can compromise plant growth and yield (63).

Some fungal strains showed promising results, suggesting their potential use in enhancing various physiological parameters. Juveniles inoculated with *Phoma* sp. recovered better after drought stress, as indicated by higher Ψ_L_ following the rewatering period. The plants also showed improved leaf recovery after the rewatering period. This phenomenon can be attributed to improved regulation of stomatal opening and reduced water loss through transpiration during the recovery phase (60, 62). The inoculated juveniles can maintain a more favorable osmotic balance, characterized by a lower solute concentration and higher water content, compared to non-inoculated plants or those inoculated with other fungi (64). In addition, a previous study showed that *Phoma* sp., when inoculated on *Pinus tabulaeformis*, can induce drought tolerance by significantly enhancing antioxidant activity (54).

A decrease in Ks was observed on days 23 and 34 in the non-drought control plants compared to day 0. This decline may be attributed to the significant temperature increase during this period. Elevated temperatures can trigger stomatal closure as a protective mechanism to reduce water loss and prevent the accumulation of reactive oxygen species (ROS) (65). It is important to note that, although stomatal closure occurred, the observed values remained within the optimal range reported for *T. cacao* (66).

Proline is an amino acid produced in response to water stress. It plays several critical roles in plants, including protection, maintaining osmotic balance, stabilizing membranes, and eliminating reactive oxygen species (ROS) (9, 55). Proline accumulates in plants in response to abiotic stressors, such as drought, protecting against these stresses (55, 67). Proline acts as a protective mechanism by stabilizing the membrane and neutralizing reactive oxygen species (ROS), which affects the plant’s water potential (68, 69). Our results showed that *T. cacao* ICS95 juveniles exposed to drought conditions experienced a significant increase in proline production compared to non-drought control plants. The highest concentrations of proline were observed in non-inoculated juveniles and those inoculated with *Acrophialophora* sp. under drought conditions. Previous reports have shown higher abundances of this genus in saline soils (70), in tropical soils, and it is considered a thermotolerant genus (57). However, to the best of our knowledge, it has not been reported to affect proline content in plants under drought stress.

In contrast, plants inoculated with *Phoma* sp. *Ectophoma sp.* and *Didymocrea* sp., exposed to drought, exhibited significantly smaller increases in proline levels than those under normal conditions. This result is intriguing because, according to Zhou et al. *Phoma* spp. can increase proline levels in plants such as *Pinus tabulaeformis* under drought conditions (54). This increase is associated with the modulation of abscisic acid (ABA) signaling pathways, as ABA concentrations rise in response to low water availability. Elevated ABA levels promote the production of molecules associated with the induction of closing stomata as a protective response to water stress (71). Such a discrepancy was also observed with Arbuscular mycorrhizal (AM) fungi (72). While drought stress induced an accumulation of proline in several AM plants, in other species, the concentration of proline in shoots was lower (72). The authors concluded that AM plants are less affected by water stress, resulting in lower proline levels due to the simultaneous inhibition of their synthesis and enhancement of their degradation (72). More studies are needed to determine if a similar phenomenon occurs in our research when cacao plants are inoculated with *Phoma* spp. and other genera that showed lower proline levels. A key finding for future studies is that cacao juveniles inoculated with *Phoma* sp. also exhibited lower proline accumulation in normal, non-drought conditions, suggesting that the fungus alters the proline metabolism.

Drought is considered one of the most significant abiotic stresses affecting the productivity and growth of *T. cacao* (73). According to Verbraeken et al. plants experience a reduction in growth in response to water stress, as the limited availability of water hinders nutrient absorption, which is essential for plant development (74). Additionally, the loss of cellular turgor further compromises critical cellular processes required for growth (58, 75). These factors help explain the significant reduction in RGR observed in both inoculated and non-inoculated juvenile plants exposed to drought compared to those grown under non-drought stress conditions.

Finally, it has been reported that endophytic fungi can promote plant growth and increase leaf area through mechanisms such as the stimulation of phytohormones (76). This could explain the significant increase in leaf area and RGR observed in *T. cacao* juveniles inoculated with endophytic fungi, particularly with *Phoma* sp., previously reported as a growth promoter (54). A similar increase in leaf area around new leaves was observed in juveniles inoculated with *Acrophialophora* sp., *Ectophoma sp.*, and *Fusarium* sp. the latter showed the most remarkable increase in leaf area. *Ectophoma sp.* has not been previously reported with such growth-promoting effects, in contrast to *Acrophialophora* sp. and *Fusarium* sp., which have been recognized as growth promoters in plants such as tomato, cucumber, and *Arabidopsis thaliana* (57, 77, 78).

This study provides the first comprehensive analysis of endophytic fungi associated with the cactus *Stenocereus* spp. and evaluates their potential to enhance drought tolerance in vitro. Our findings reveal that both Tatacoa and Taganga display high diversity indices, reflecting a wide variety of species and complex community structures essential for maintaining ecosystem stability and resilience. Furthermore, the fungal root endophytes of *Stenocereus* spp. demonstrated significant potential as growth promoters for juvenile *Theobroma cacao* ICS95 plants, suggesting their applicability in sustainable agricultural practices. This work contributes valuable insights to developing biotechnological strategies to mitigate the adverse effects of climate change, particularly in drought-affected regions. The results support using fungal endophytes as a promising tool for enhancing agricultural resilience and sustainability in increasingly variable environmental conditions.

## Funding sources and collecting permits

This research was funded by a Seed Project award number INV-2023-174-2915 from the School of Sciences at Universidad de los Andes, and some funding from the Sequencing Core Facility (Gencore) of the same institution. The root samples of the present research were collected under the collection permit number 20236200291612 of ANLA.

## Declaration of generative AI and AI-assisted technologies in the writing process

We used ChatGPT and Grammarly AI tools to improve the document’s quality, grammar, and coherence, without altering the core concepts or introducing new ideas. After using this tool/service, the authors reviewed and edited the content as needed and take full responsibility for the content of the published article.

## Author contributions

KST-O wrote the original draft and conducted the experiments. AM-S, FR, and MG-S contributed to the manuscript’s conceptualization, methodology, and editing. SR ensured the funding, supervised the work, and edited the manuscript.

## Supplementary Material

**Supplementary Table 1.** Geographic information of localities where the roots of *<u>Stenocereus</u>* spp. were sampled.

| Department | Locality | Sample | Elevation<br>(m.a.s.l) | Latitude | Longitude |
| --- | --- | --- | --- | --- | --- |
| Huila | Tatacoa | 1 | 489 | 3.227898 | -75.150175 |
| Huila | Tatacoa | 2 | 469 | 3.228118 | -75.15031 |
| Huila | Tatacoa | 3 | 470 | 3.228342 | -75.150432 |
| Huila | Tatacoa | 4 | 469 | 3.228132 | -75.149673 |
| Huila | Tatacoa | 5 | 473 | 3.228327 | -75.149527 |
| Huila | Tatacoa | 6 | 466 | 3.228808 | -75.150205 |
| Huila | Tatacoa | 7 | 466 | 3.228785 | -75.150878 |
| Huila | Tatacoa | 8 | 466 | 3.229205 | -75.1512 |
| Huila | Tatacoa | 9 | 469 | 3.228932 | -75.150497 |
| Huila | Tatacoa | 10 | 466 | 3.228808 | -75.150205 |
| Huila | Tatacoa | 11 | 466 | 3.229547 | -75.150115 |
| Huila | Tatacoa | 12 | 464 | 3.229375 | -75.149462 |
| Magdalena | Taganga | 1 | 19 | 11.268545 | -74.194403 |
| Magdalena | Taganga | 2 | 18 | 11.268445 | -74.194498 |
| Magdalena | Taganga | 3 | 21 | 11.26837 | -74.194575 |
| Magdalena | Taganga | 4 | 32 | 11.26854 | -74.194612 |
| Magdalena | Taganga | 5 | 29 | 11.268662 | -74.194668 |
| Magdalena | Taganga | 6 | 34 | 11.268818 | -74.19467 |
| Magdalena | Taganga | 7 | 33 | 11.268642 | -74.194708 |
| Magdalena | Taganga | 8 | 21 | 11.268735 | -74.195045 |
| Magdalena | Taganga | 9 | 14 | 11.259083 | -74.194953 |
| Magdalena | Taganga | 10 | 14 | 11.26913 | -74.195888 |
| Magdalena | Taganga | 11 | 30 | 11.26905 | -74.196407 |
| Magdalena | Taganga | 12 | 24 | 11.269178 | -74.19608 |

**Supplementary Table 2.** Morphological and molecular identification of the fungal isolates.

| Best Blastn Result with ITS primers | Best Blastn Result with TEF primers | NCBI ID | Microscopic description | Macroscopic description | Morphology Identification | Locality |
| --- | --- | --- | --- | --- | --- | --- |
| *Acrocalymma* sp. |  | PV938402 | No data available. | Velvety colony with white to grey color (approx. RGB 255, 242, 206 ) | No data available. | Tatacoa (Huila) |
| *Acrophialophora* sp. |  | PX218687 | Dark, thick, septate mycelium. Bottle-shaped phialides producing hyaline, ellipsoidal ameroconidia. Pycnidia formation observed. | Colony with leathery to velvety texture and grey color (approx. RGB 0, 72, 66). White ring at the outer margin (approx. RGB 254, 249, 255) | *Acrophialophora* sp. | Taganga (Magdalena) |
| *Alternaria* sp. |  | PV938419 | Mycelium is dematiaceous, thick-walled, and septate. Dictyoconidia dematiaceous produced in acropetal chains. | Velvety brown colony in the anverse and reverse (approx. RGB 134, 93, 0). | *Alternaria* sp. | Tatacoa (Huila) - Taganga (Magdalena) |
| *Aspergillus arcoverdensis* |  | PV9384<br>25 | Hyaline, thin mycelium. Conidiophores with terminal vesicles bearing bottle-shaped phialides, from which acropetal chains of hyaline, ovoid conidia arise. | Colony granular with green color in the anverse (approx. RGB 134, 188, 0) and white in the reverse (approx. RGB 252, 255, 226). | *Aspergillus* sp. | Tatacoa (Huila) - Taganga (Magdalena) |
| *Aspergillus eucalypticola* |  | PV9383<br>89 | Hyaline, thin mycelium. Conidiophores with terminal vesicles bearing bottle-shaped phialides, from which acropetal chains of hyaline, ovoid conidia arise. | Colony with a granular texture, black on the surface (approx. RGB 0, 41, 4) and white (approx. RGB 252, 255, 226) on the reverse side. | *Aspergillus* sp. | Tatacoa (Huila) - Taganga (Magdalena) |
| *Aspergillus flavus* |  | PV9384<br>12 | Hyaline, thin mycelium. Conidiophores with terminal vesicles bearing bottle-shaped phialides, from which acropetal chains of hyaline, ovoid conidia arise. | Colony with granular texture, green to yellow color (approx. RGB 203, 200, 3) in the anverse and grey in the reverse (approx. RGB 0, 84, 85). | *Aspergillus* sp. | Tatacoa (Huila) - Taganga (Magdalena) |
| *Aspergillus heldtiae* |  | PV9384<br>08 | Hyaline, thin mycelium. Conidiophores with terminal vesicles bearing bottle-shaped phialides, from which acropetal chains of hyaline, ovoid conidia arise. | Colony with granular texture, yellow (approx. RGB 254, 232, 38) color in the anverse and yellow-green (approx. RGB 196, 213, 0) in the reverse. | *Aspergillus* sp. | Tatacoa (Huila) - Taganga (Magdalena) |
| *Aspergillus lentulus* |  | PV9384<br>26 | Hyaline, thin mycelium. Conidiophores with terminal vesicles bearing bottle-shaped phialides, from which acropetal chains of hyaline, ovoid conidia arise. | Velvety colony white with pink in the center (approx. RGB 250, 237, 255). | *Aspergillus* sp. | Tatacoa (Huila) - Taganga (Magdalena) |
| *Aspergillus neoterreus* |  | PV9384<br>03 | Hyaline, thin mycelium. Conidiophores with terminal vesicles bearing bottle-shaped phialides, from which acropetal chains of hyaline, ovoid conidia arise. | Colony with a granular texture, black on the surface (approx. RGB 0, 70, 70) and white (approx. RGB 255, 239, 229) on the reverse side. | *Aspergillus* sp. | Tatacoa (Huila) - Taganga (Magdalena) |
| *Aspergillus niger* |  | PV9384<br>20 | Hyaline, thin mycelium. Conidiophores with terminal vesicles bearing bottle-shaped phialides, from which acropetal chains of hyaline, ovoid conidia arise. | Colony granular with black color (approx. RGB 2, 26, 0) in the anverse and white (approx. RGB 248, 255, 250) in the reverse. | *Aspergillus* sp. | Tatacoa (Huila) - Taganga (Magdalena) |
| *Aspergillus terreus* |  | PV9384<br>18 | Hyaline, thin mycelium. Conidiophores with terminal vesicles bearing bottle-shaped phialides, from which acropetal chains of hyaline, ovoid conidia arise. | Colony granular with yellow color (approx. RGB 245, 215, 41) in the anverse and brown (approx. RGB 178, 118, 1) in the reverse. In some cases, yellow synnemata are formed. | *Aspergillus* sp. | Tatacoa (Huila) - Taganga (Magdalena) |
| *Aureobasidium* sp. |  | PV9383<br>95 | No data available. | Velvety white to grey (approx. RGB 244, 230, 255) in the anverse and grey (approx. RGB 1, 68, 86) in the reverse. | No data available. | Tatacoa (Huila) - Taganga (Magdalena) |
| *Clavulina* sp. |  | PV9384<br>05 | No data available. | Velvety colony with grey to pink color (approx. RGB 254, 221, 237). | No data available. | Tatacoa (Huila) |
| *Coprinellus* sp. |  | PV9384<br>09 | No data available. | Velvety colony with white color (approx. RGB 246, 254, 243) in the anverse and pink to borwn (approx. RGB255, 192, 203) in the reverse. | No data available. | Tatacoa (Huila) |
| *Curvularia alcornii* |  | PV9384<br>10 | Mycelium is dematiaceous, thick-walled, and septate. Fragmaconidia dematiaceous produced from sympodial conidiogenous cells. | Velvety colony with grey color (approx. RGB 0, 92, 71) in the anverse and black color (approx. RGB 217, 30, 0) in the reverse. | *Curvularia* sp. | Tatacoa (Huila) - Taganga (Magdalena) |
| *Curvularia buchloes* |  | PV9384<br>01 | Mycelium is dematiaceous, thick-walled, and septate. Fragmaconidia dematiaceous produced from sympodial conidiogenous cells. | Velvety grey to black (approx. RGB 1, 14, 58) in the anverse and in the reverse | *Curvularia* sp. | Tatacoa (Huila) - Taganga (Magdalena) |
| *Curvularia coimbatorensis* |  | PV9383<br>93 | Mycelium is dematiaceous, thick-walled, and septate. Fragmaconidia dematiaceous produced from sympodial conidiogenous cells. | Velvety colony with grey color (approx. RGB 0, 92, 71). The reverse was black (approx. RGB 217, 30, 0). | *Curvularia* sp. | Tatacoa (Huila) - Taganga (Magdalena) |
| *Curvularia milisiae* |  | PV9384<br>07 | Mycelium is dematiaceous, thick-walled, and septate. Fragmaconidia dematiaceous produced from sympodial conidiogenous cells. | Velvety black (approx. RGB 0, 27, 24) in the anverse and in the reverse. | *Curvularia* sp. | Tatacoa (Huila) - Taganga (Magdalena) |
| *Curvularia pseudolunata* |  | PV9383<br>94 | Mycelium is dematiaceous, thick-walled, and septate. Fragmaconidia dematiaceous produced from sympodial conidiogenous cells. | Velvety colony brown (approx. RGB 78, 64, 0) with orange lines (approx. RGB 174, 95, 0) and reverse orange (approx. RGB 174, 95, 0) with black lines (approx. RGB 44, 34, 0). | *Curvularia* sp. | Tatacoa (Huila) - Taganga (Magdalena) |
| *Curvularia simmonsii* |  | PV9384<br>00 | Mycelium is dematiaceous, thick-walled, and septate. Fragmaconidia dematiaceous produced from sympodial conidiogenous cells. | Velvety colony grey (approx. RGB 30, 46, 1) in the anverse and in the reverse. | *Curvularia* sp. | Tatacoa (Huila) - Taganga (Magdalena) |
| *Deniquelata* sp. |  | PV9384<br>27 | Mycelium is hyaline, thin, and sparsely septate. Subglobose conidia produces by conidiogenous cells subcylindrical. | Velvety colony white in the anverse and in the reverse (approx. RGB 246, 254, 231), with ring at the outer margin yellow (approx. RGB 255, 234, 166). | No data available. | Tatacoa (Huila) |
| *Diaporthe* sp. |  | PV9384<br>22 | Mycelium is hyaline, thin, and sparsely septate. Ameroconidia fusiform. | Velvety-Cottony white (approx. RGB 246, 254, 231) colony in the anverse and in the reverse with ring at the outer margin black. | No data available. | Tatacoa (Huila) |
| *Didymella* sp. |  | PV9384<br>13 | Mycelium is hyaline, thin, and sparsely septate. No reproductive structures were observed under the examined conditions. | Velvety grey (approx. RGB 30, 46, 1) colony in the anverse and in the reverse with ring at the outer margin white (approx. RGB 246, 254, 231). | No data available. | Tatacoa (Huila) |
| *Didymella* sp. |  | PV9384<br>24 | Mycelium is hyaline, thin, and sparsely septate. No reproductive structures were observed under the examined conditions. | Colony cottony white (approx. RGB 246, 254, 231) in the anverse and in the reverse. | No data available. | Tatacoa (Huila) |
| *Didymocrea* sp. |  | PX2186<br>85 | Dark, thick, and septate mycelium. Micronematous conidiophores producing solitary, dematiaceous conidia acrogenously. | Velvety colony with grey (approx. RGB 30, 46, 1) color in the anverse and black color in the reverse (approx. RGB 0, 8, 37). | *Didymocrea* sp. | Tatacoa (Huila) |
| *Ectophoma* sp. |  | PX2186<br>86 | Dark, septate mycelium and produced brown pycnidia containing ellipsoidal ameroconidia. Each conidium presented two guttules, and chlamydospores were observed forming in chains | Velvety-leathery colony with grey (approx. RGB 30, 46, 1) color in the anverse and black color in the reverse (approx. RGB 0, 8, 37). | *Phoma* sp. | Tatacoa (Huila) - Taganga (Magdalena) |
| *Epicoccum* sp. |  | PV9383<br>96 | No data available. | Colony with leathery texture and black color (approx. RGB 0, 8, 37). | No data available. | Tatacoa (Huila) - Taganga (Magdalena) |
| *Fusarium falciforme* | *Neocosmospora rubicola* | FEC28_1507202<br>5 | Hyaline, thin, septate mycelium. Ameroconidia and hyaline macroconidia (didymoconidium), produced on flask-shaped phialides. | Cottony, ivory white in color (approx. RGB 254, 243, 213), with scarce aerial mycelium. The colony surface was smooth and uniform, with entire margins. | *Fusarium* sp. | Tatacoa (Huila) - Taganga (Magdalena) |
| *Fusarium*<br>*falciforme* | *Neocosmos*<br>*pora* sp. | FEC30_<br>1507202<br>5 | Hyaline, thin, septate<br>mycelium.<br>Ameroconidia and<br>hyaline macroconidia<br>(didymoconidium),<br>produced on flask-shaped phialides. | Cottony, ivory white<br>in color (approx. RGB<br>254, 243, 213), with<br>scarce aerial<br>mycelium. The<br>colony surface was<br>smooth and uniform,<br>with entire margins. | *Fusarium* sp. | Tatacoa (Huila) - Taganga<br>(Magdalena) |
| *Fusarium*<br>*falciforme* | *Neocosmos*<br>*pora* sp. | FEC31_<br>1507202<br>5 | Hyaline, thin, septate<br>mycelium.<br>Ameroconidia and<br>hyaline macroconidia<br>(didymoconidium),<br>produced on flask-shaped phialides. | Cottony, ivory white<br>in color (approx. RGB<br>254, 243, 213), with<br>scarce aerial<br>mycelium. The<br>colony surface was<br>smooth and uniform,<br>with entire margins. | *Fusarium* sp. | Tatacoa (Huila) - Taganga<br>(Magdalena) |
| *Fusarium* sp. | *Fusarium*<br>sp. | FEC29_<br>1507202<br>5 | Hyaline, thin, septate<br>mycelium. Hyaline<br>didymoconidium micro<br>and macroconida,<br>produced on flask-shaped phialides. | Cottony, ivory white<br>in color (approx. RGB<br>254, 243, 213), with<br>scarce aerial<br>mycelium. The<br>colony surface was<br>smooth and uniform,<br>with entire margins. | No data<br>available. | Tatacoa (Huila) - Taganga<br>(Magdalena) |
| *Lasiodiplodia* sp. |  | PV9384<br>21 | Mycelium is septate<br>and hyaline. No<br>reproduction structures | Colony cottony grey<br>(approx. RGB 1, 22,<br>51) in the anverse and<br>black in the reverse<br>(approx. RGB 0, 8,<br>37). | *Lasiodiplodia*<br>sp. | Tatacoa (Huila) - Taganga<br>(Magdalena) |
| *Latorua* sp. |  | PV9383<br>91 | Hyaline, thin, septate mycelium. Simple, geniculate, macronematous conidiophore. Ellipsoidal conidia in acropetal chains. | Velvety colony with grey color (approx. RGB 1, 22, 51) with ring at the outer margin. The reverse was grey. | *Acroconidiella* sp. | Taganga (Magdalena) |
| *Macrophomina* sp. |  | PV9384<br>15 | No data available. | Velvety colony grey (approx. RGB 1, 22, 51) in the anverse and in the reverse. | *Macrophomina* sp. | Tatacoa (Huila) - Taganga (Magdalena) |
| *Moniliophthora* sp. |  | PV9384<br>14 | No data available. | Velvety white (approx. RGB 1, 22, 51) in the anverse and in the reverse | No data available. | Tatacoa (Huila) - Taganga (Magdalena) |
| *Monosporascus* sp. |  | PV9383<br>92 | No data available. | Cottony, ivory white in color (approx. RGB 242, 254, 232), with scarce aerial mycelium. The colony surface was smooth and uniform, with entire margins. | No data available. | Tatacoa (Huila) - Taganga (Magdalena) |
| *Myrmaecium* sp. |  | PV9384<br>23 | No data available. | Velvety grey colony (approx. RGB 1, 22, 51) in the anverse and brown (approx. RGB 139, 117, 0) in the reverse. | No data available. | Taganga (Magdalena) |
| *Oblongocollomyces zhivanae* |  | PV9384<br>04 | No data available. | Velvety colony with grey color (approx. RGB 20, 44, 66). | No data available. | Tatacoa (Huila) |
| *Paraboeremia* sp. |  | PV9383<br>98 | No data available. | Colony with leathery texture and brown color (approx. RGB 94, 33, 1). | No data available. | Tatacoa (Huila) |
| *Phoma* sp. |  | PX2186<br>88 | Mycelium is thick-walled and septate, ranging from hyaline to dematiaceous. Pycnidia are dark, globose to subglobose, and ostiolate. Conidia formed within pycnidia are ovoid, aseptate, and vary in pigmentation from hyaline to dematiaceous. Each conodium with two guttules. | Velvety colony with brown-yellow (approx. RGB 124, 116, 2) color in the anverse. | *Phoma* sp. | Tatacoa (Huila) |
| *Poaceascoma* sp. |  | PV9384<br>11 | No data available. | Velvety colony with brown-grey color in the anverse and black color in the reverse (approx. RGB 87, 58, 0). | No data available. | Tatacoa (Huila) - Taganga (Magdalena) |
| *Polyporus* sp. |  | PV9383<br>90 | Mycelium hyaline, thin, and sparsely septate. No reproductive structures were observed. | The colony was white (approx. RGB 254, 243, 213) with a yellow (approx RGB 255, 245, 196) ring at the outer margin and a velvety texture on the surface. The reverse was initially white (approx. RGB 254, 243, 213), becoming brown (approx. RGB 223, 168, 1) at the center over time. | No data available. | Taganga (Magdalena) |
| *Purpureocillium lilacinum* |  | PV9384<br>06 | Hyaline, septate mycelium with erect conidiophores bearing flask-shaped phialides that produce chains of hyaline, ellipsoidal conidia. | Velvety colony with brown color (approx. RGB 97, 34, 1) in anverse and reverse. Production of yellow pigment (approx. RGB 193, 164, 1). | *Purpureocillium* sp. | Tatacoa (Huila) |
| *Rhizopus sexualis* |  | PV9383<br>99 | Dark, thick, and sparsely septate mycelium. Presence of dematiaceous sporangia and sporangiophores, along with well-developed rhizoids. | Cotony cottony white-grey (approx. RGB 237, 245, 254) in the anverse and grey in the reverse. | *Rhizopus* sp. | Tatacoa (Huila) - Taganga (Magdalena) |
| *Rhodotorula* sp. |  | PV9384<br>17 | Cells are unicellular, globose to ovoid, and reproduce by multilateral budding. | Colonies are smooth, moist to mucoid, and display a characteristic pink color (approx. RGB 255, 174, 237). | *Rhodotorula* sp. | Taganga (Magdalena) |
| *Trichoderma saturnisporum* |  | PV9383<br>97 | Septate, hyaline mycelium with branched conidiophores bearing flask-shaped phialides that produce blue ameroconidia. | Cottony texture with white color (approx. RGB 236, 255, 230) with coconcentric green-blue rings (approx. RGB 1, 138, 58). | *Trichoderma* sp. | Tatacoa (Huila) - Taganga (Magdalena) |
| *Trichoderma* sp. |  | PV9384<br>16 | Septate, hyaline mycelium with branched conidiophores bearing flask-shaped phialides that produce green ameroconidia. | Colony cottony white (approx. RGB 253, 255, 245) in the anverse and in the reverse. | *Trichoderma* sp. | Taganga (Magdalena) |
| *Cylindrocarpon* sp | . | FEC32_1507202<br>5 | Hyaline, thin, septate mycelium. Hyaline didymoconidium micro and macroconida, produced on flask-shaped phialides. | Cottony, ivory white in color (approx. RGB 254, 243, 213), with scarce aerial mycelium. The colony surface was smooth and uniform, with entire margins. | *Fusarium* sp. | Tatacoa (Huila) - Taganga (Magdalena) |
| *Bisifusarium* sp. |  | FEC49_1507202<br>5 | Hyaline, thin, septate mycelium. Hyaline didymoconidium micro and macroconida, produced on flask-shaped phialides. | Cottony, ivory white in color (approx. RGB 251, 240, 254), with scarce aerial mycelium. The colony surface was smooth and uniform, with entire margins. | *Fusarium* sp. | Tatacoa (Huila) - Taganga (Magdalena) |

**Supplementary Table 3.**
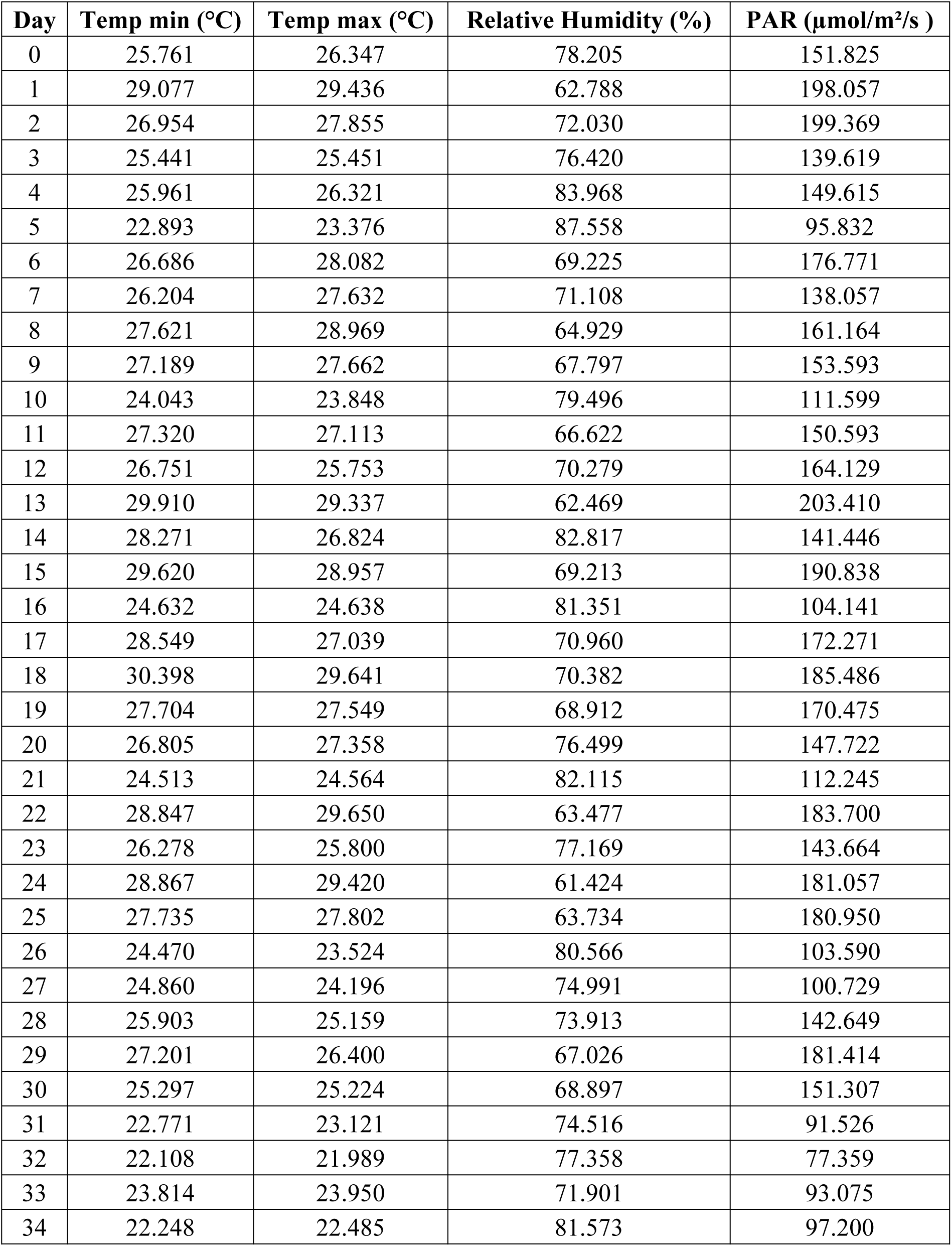
Daily (7:00 am – 6:00 pm) average environmental conditions during the greenhouse assays.

| Day | Temp min (°C) | Temp max (°C) | Relative Humidity (%) | PAR (μmol/m<sup>2</sup>/s ) |
| --- | --- | --- | --- | --- |
| 0 | 25.761 | 26.347 | 78.205 | 151.825 |
| 1 | 29.077 | 29.436 | 62.788 | 198.057 |
| 2 | 26.954 | 27.855 | 72.030 | 199.369 |
| 3 | 25.441 | 25.451 | 76.420 | 139.619 |
| 4 | 25.961 | 26.321 | 83.968 | 149.615 |
| 5 | 22.893 | 23.376 | 87.558 | 95.832 |
| 6 | 26.686 | 28.082 | 69.225 | 176.771 |
| 7 | 26.204 | 27.632 | 71.108 | 138.057 |
| 8 | 27.621 | 28.969 | 64.929 | 161.164 |
| 9 | 27.189 | 27.662 | 67.797 | 153.593 |
| 10 | 24.043 | 23.848 | 79.496 | 111.599 |
| 11 | 27.320 | 27.113 | 66.622 | 150.593 |
| 12 | 26.751 | 25.753 | 70.279 | 164.129 |
| 13 | 29.910 | 29.337 | 62.469 | 203.410 |
| 14 | 28.271 | 26.824 | 82.817 | 141.446 |
| 15 | 29.620 | 28.957 | 69.213 | 190.838 |
| 16 | 24.632 | 24.638 | 81.351 | 104.141 |
| 17 | 28.549 | 27.039 | 70.960 | 172.271 |
| 18 | 30.398 | 29.641 | 70.382 | 185.486 |
| 19 | 27.704 | 27.549 | 68.912 | 170.475 |
| 20 | 26.805 | 27.358 | 76.499 | 147.722 |
| 21 | 24.513 | 24.564 | 82.115 | 112.245 |
| 22 | 28.847 | 29.650 | 63.477 | 183.700 |
| 23 | 26.278 | 25.800 | 77.169 | 143.664 |
| 24 | 28.867 | 29.420 | 61.424 | 181.057 |
| 25 | 27.735 | 27.802 | 63.734 | 180.950 |
| 26 | 24.470 | 23.524 | 80.566 | 103.590 |
| 27 | 24.860 | 24.196 | 74.991 | 100.729 |
| 28 | 25.903 | 25.159 | 73.913 | 142.649 |
| 29 | 27.201 | 26.400 | 67.026 | 181.414 |
| 30 | 25.297 | 25.224 | 68.897 | 151.307 |
| 31 | 22.771 | 23.121 | 74.516 | 91.526 |
| 32 | 22.108 | 21.989 | 77.358 | 77.359 |
| 33 | 23.814 | 23.950 | 71.901 | 93.075 |
| 34 | 22.248 | 22.485 | 81.573 | 97.200 |

**Supplementary Table 4.**
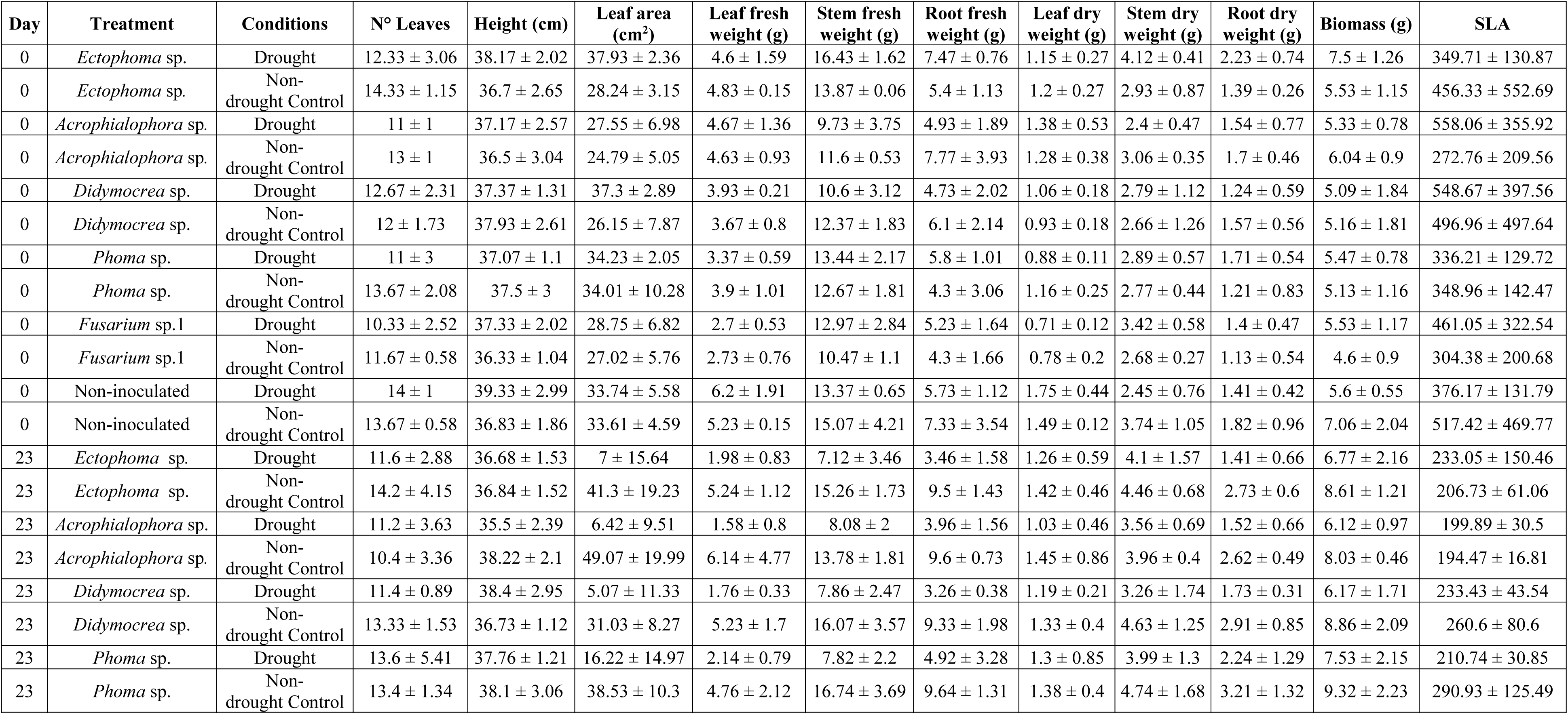
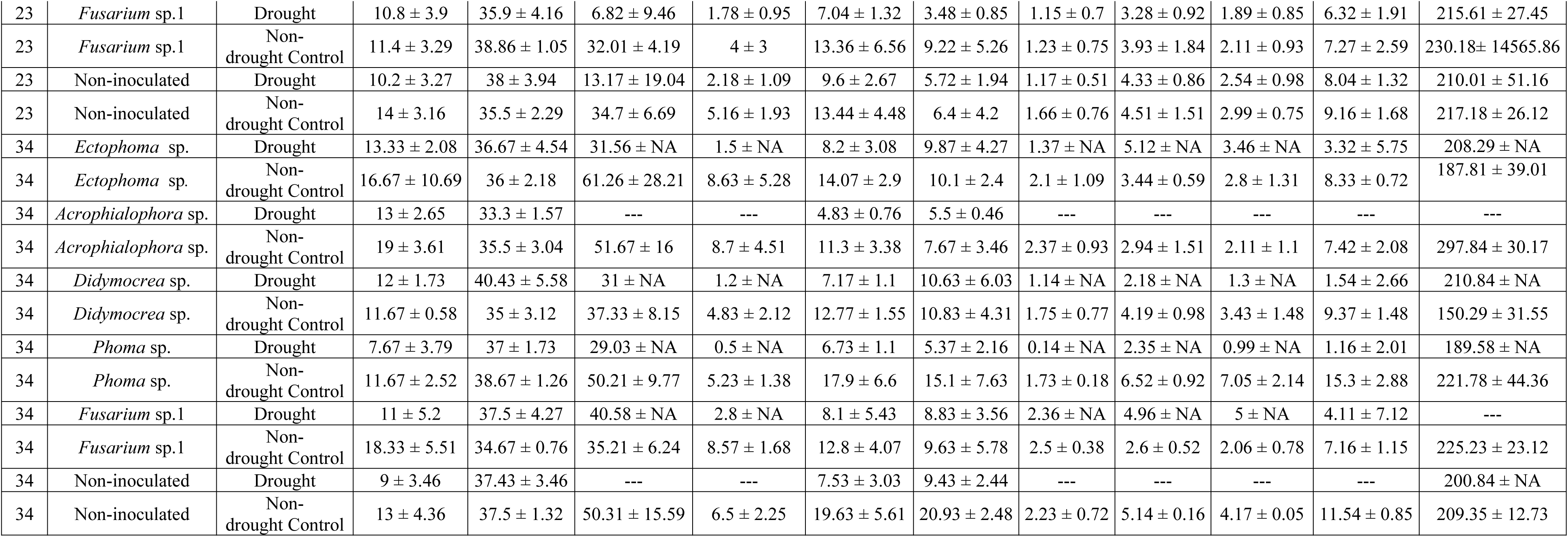
Means of morphological parameters of juveniles of Theobroma cacao ICS95 exposed under drought and normal conditions, measured on different days of the experiment.

**Supplementary Figure 1.**
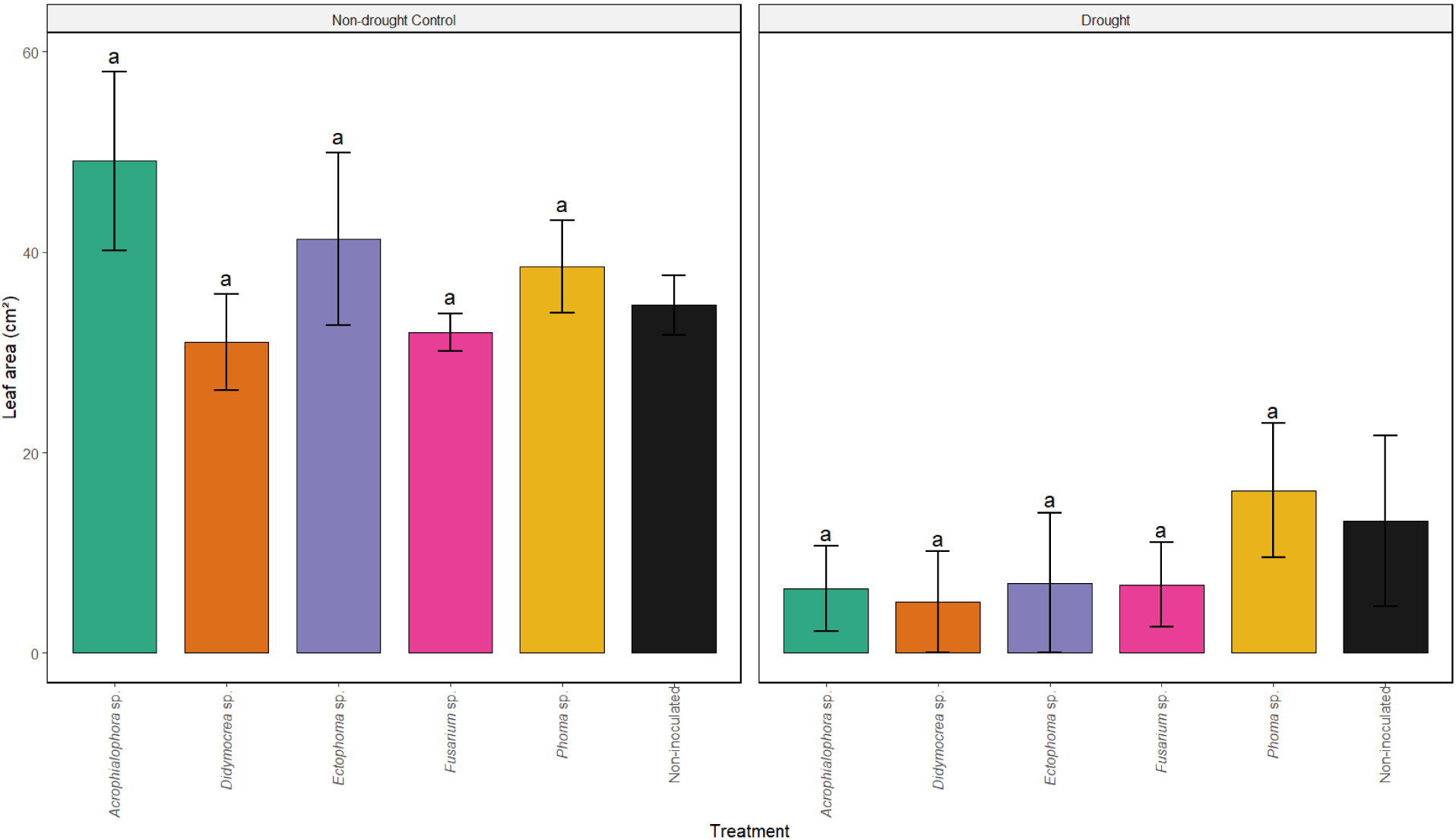
Mean leaf area (cm²) of *Theobroma cacao* ICS95 juveniles exposed to non-drought control (left) and drought conditions for 13 days (right) (day 23).

**Supplementary Figure 2.**
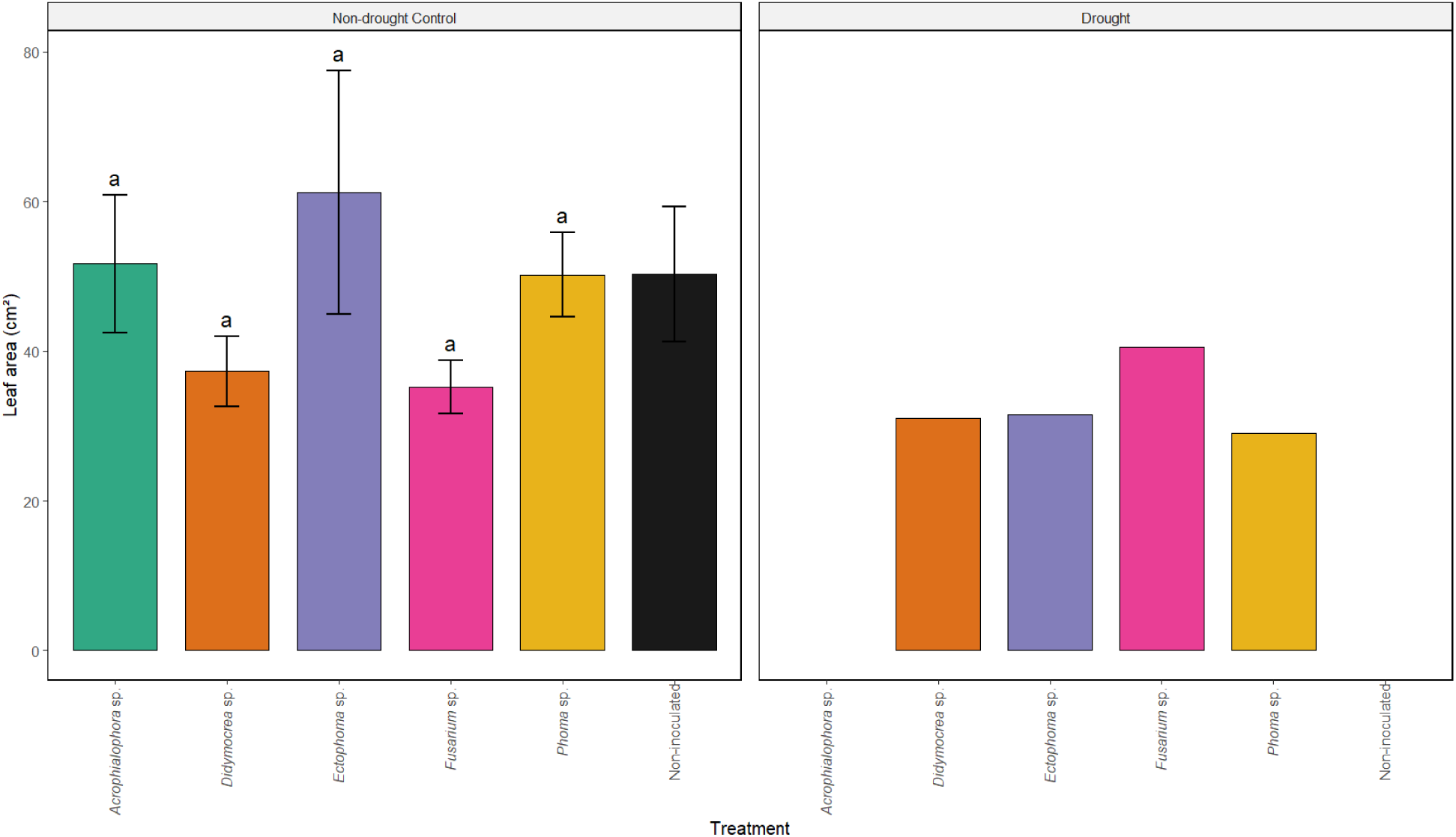
Mean leaf area (cm²) of *Theobroma cacao* ICS95 juveniles exposed to non-drought control (left) and drought conditions followed by rewatering for two weeks (right). The figure does not include error bars because at this stage, the plants showed very few leaves.

